# Comprehensive Survey of Gut Microbiome Associations with Health Conditions in the Human Phenotype Project

**DOI:** 10.1101/2025.06.26.661672

**Authors:** Daphna Weissglas-Volkov, Yotam Reisner, Tal Shor, Alon Diament, Adam Jankelow, Anastasia Godneva, Brooke A Napier, Raja Dhir, Eran Segal

## Abstract

The human gut microbiome is increasingly implicated with diverse health conditions, highlighting the importance of developing comprehensive resources to systematically map these associations. In this study, we leveraged the Human Phenotype Project (HPP) 10K cohort, a large, deeply phenotyped population study cohort with extensive metagenomic profiling, to explore associations between the gut microbiome and 37 health indications. These include conditions with established or emerging links to the microbiome, such as obesity, diabetes, hyperlipidemia, metabolic syndrome, inflammation, liver disease, cardiovascular and kidney conditions, immune and allergic diseases, gastrointestinal symptoms, mental health, and sleep-related traits. Using a curated set of approximately 200 refined phenotypic features and analyzing 1,184 microbial species, we performed robust statistical association analyses. We observed significant enrichment of associations in 14 health indications, with most associations reflecting increased microbial abundance in favorable health states, thereby suggesting potential microbial targets for intervention. This work introduces a publicly available, high-resolution resource to facilitate future research and support the development of microbiome-informed health strategies.

## Introduction

The human gut microbiome is a complex and diverse ecosystem made up of trillions of microorganisms, including bacteria, archaea, viruses, and fungi, residing in the gastrointestinal tract [1]. These microorganisms play crucial roles in human health by aiding digestion, producing essential nutrients, modulating immune responses, and protecting against harmful pathogens [2]. There is strong evidence linking gut bacteria to various health conditions [3,4]. For example, digestive disorders like inflammatory bowel disease (IBD) and irritable bowel syndrome (IBS) are linked to imbalances in gut microbes, which worsen inflammation and symptoms [5]. Metabolic disorders, such as obesity and type 2 diabetes, are associated with the microbiome, as it affects energy metabolism, insulin sensitivity, and glucose regulation [6, 7]. Furthermore, cardiovascular diseases are influenced by microbial metabolites like trimethylamine-N-oxide (TMAO), which promote atherosclerosis [8]. These examples reflect the vital role of the gut microbiome in supporting metabolic and inflammatory equilibrium [3].

Emerging research suggests that the gut microbiome may also affect a broader range of health conditions. Mental health disorders, such as depression and anxiety, have been associated with gut bacteria through the gut-brain-axis, which impacts neurotransmitter production and inflammatory pathways [9,10]. Neurodegenerative diseases, including Parkinson’s and Alzheimer’s, are being explored for their potential links to gut microbiota [11,12]. Likewise, autoimmune diseases like multiple sclerosis and rheumatoid arthritis may result from microbial effects on immune system regulation [13, 14]. Traits such as ADHD are also under investigation for possible microbiome involvement [15]. These research underscore an increasing interest in how gut bacteria could influence systemic and neural or cognitive health [16,17].

The connection between gut bacteria and various health conditions emphasizes the importance of maintaining a balanced microbiome. Probiotics, live beneficial microorganisms, are becoming increasingly popular for their ability to support or restore gut health. Despite the progress in microbiome research, significant gaps remain in understanding which bacterial strains are most beneficial and their mechanism of action [18]. One of the reasons is the lack of consistency in methodologies that creates barriers to identifying the specific bacteria involved. Comparing findings across studies is challenging due to variability in study designs, sample populations, environmental factors, bacterial taxa examined, and the health traits analyzed [19]. Large-scale population-based cohort studies with comprehensive and detailed health profiling offer a promising opportunity for rigorous microbiome research [20]. These studies provide the statistical power necessary to detect subtle associations and enable a broad investigation into the various interactions between microbiome and health traits using the same study design and analytical approach. Establishing standardized approaches could facilitate meaningful comparisons of findings across traits and microbial species.

The Human Phenotype Project (HPP) 10K cohort is an extensive population-based research initiative that integrates in-depth health-related data and longitudinal profiling with metagenomic profiles from a large group of participants [21,22]. In this study, we selected 37 health indications available within the HPP cohort, spanning metabolic, inflammatory, gastrointestinal, immune, neurological, musculoskeletal, dermatological, and sleep-related domains. Although not exhaustive, this range reflects many conditions with established or emerging microbiome links and offers a broad view of microbiome-health associations. By utilizing this uniquely rich dataset, this work aims to identify associations between the microbiome and health advancements. Furthermore, it aims to conduct a systematic exploration of a wide array of microbial species and traits using a substantial robust sample size, offering valuable insights that could guide the development of more effective probiotics and therapeutic strategies, and will be made publicly available to serve the broader scientific community.

## Methods

### Description of cohort

This study used data from the ongoing Human Phenotype Project (HPP) 10K study. For a detailed design, see reference [21,22]. In brief, the HPP study is a large-scale prospective longitudinal cohort and biobank established in Israel in 2018, tracking a group of 40-70-year-olds. Originally aimed at recruiting 10,000 individuals, the study was named “10K cohort”. Since the study’s launch, it has grown to include 13,000 who have already completed their first visit as of December 2024 and is now called the HPP. A longitudinal follow-up is planned for a duration of 25 years, with annual questionnaires and bi-annual visits at the clinical testing center (CTC) for comprehensive profiling. Predefined medical conditions were determined as exclusions as described in [21]. Information collection was previously described [22]. In short, it includes questionnaire based data on medical history, family medical history, lifestyle, female health, nutritional habits, smoking and sleep habits. Assessment tests conducted at the CTC include anthropometric measurements, vital signs including blood pressure, complete blood count (CBC), electrocardiography (ECG), ankle–brachial pressure index (ABI), hand grip strength, imaging studies such as retinal imaging, liver and carotid ultrasound (US), dual-energy X-ray absorptiometry (DXA) scan for bone mineral density (BMD) and body composition. Molecular profiling includes human genetics, transcriptomics, gut and oral microbiome, metabolomics and immune system analysis. Continuous health monitoring measurements on the two weeks following the CTC visit include continuous glucose monitoring device in parallel to logging meals, medications, exercise in a proprietary App, and sleep monitoring by a home sleep apnea test (HSAT) device for 3 nights during the same two weeks period. Additional blood test results are obtained from participants’ electronic health records (EHRs). Blood (serum, peripheral blood mononuclear cell (PBMC) and whole blood) and stool samples are stored for future research.

### Gut Microbiome Metagenomic

Sample collection, DNA extraction, and sequencing of the 10K study participants were previously described in detail [23]. Briefly, the stool samples were collected using a standardized stool collection kit (OMNIgene·GUT OMR-200, DNA Genotek), ensuring consistent sample quality. Each participant was given a kit and was requested to collect a fecal sample at home. The collected samples were transferred at room temperature to our clinical testing center (CTC) at Weizmann Institute of Science, where they were documented and frozen at −20 °C immediately and stored at −20 °C until DNA extraction was performed. Metagenomic DNA was purified using PowerMag Microbial DNA Isolation Kit (MO BIO Laboratories,27200-4) optimized for the Tecan automated platform. Libraries for next-generation sequencing were prepared using NEBNext Ultra II DNA Library Prep Kit for Illumina (New England Biolabs, E7775) and sequenced on a NovaSeq sequencing platform (Illumina). Sequencing was performed with a 100-bp single-end reads kit and a depth of 10 million reads per sample, using Illumina unique dual sequencing indexes (IDT–Syntezza Bioscience). DNA purification, library preparation,and sequencing were performed in batches of 384 samples. A standard microbial community (ZymoBIOMICS Gut MicrobiomeStandard, D6331) was inserted into each batch for quality control. We filtered reads containing Illumina adapters and low-quality reads and trimmed low quality read edges. We detected host DNA by mapping reads to the human genome using Bowtie 2 with inclusive parameters and removed those reads. The raw sequencing data underwent stringent quality control (QC), including a minimum threshold of non-human reads (>5M), a minimum length of high-quality reads after removing adaptors (> 80bp), and the matching of DNA from stool samples with participant DNA using whole genome sequencing (WGS) human genetics sample for validation purposes. The quality-filtered metagenomic reads were mapped to our human gut microbiome reference set of >3000 high-quality species genomes [24]. The algorithm described in Leviatan *et al.* 2022 [24] was employed for taxonomic classification and abundance estimation, resulting in a microbial profile of relative abundances for all the bacteria found present in the participant gut microbiome sample.

A total of 3,209 species across 11,295 samples from 9,249 unique participants were available for this study. Missing data were assumed to represent missing abundance or abundances that are below the detection limit. Thus, missing data were imputed by a minimum value of 0.0001. We implemented additional sample QC procedures, requiring more than 4M unique mapped reads and the identification of at least 200 unique species. Resulting in 10,623 eligible samples from 8,783 participants.

To assess the diversity of the gut microbiome, we calculated several diversity metrics. Alpha diversity was measured using both the Shannon Effective Number and Simpson Reciprocal Index, which provide insights into the richness and evenness of species within each sample. Beta diversity was computed using the Bray-Curtis dissimilarity index, measuring the compositional dissimilarity between different samples. These diversity metrics were used to create a comprehensive score of gut microbiome diversity for each participant.

### Characterization of Health Indications

Based on the data available in HPP the following 37 health indications were selected for the investigation with microbiome relative abundance in alphabetic order: Attention Deficit Hyperactivity Disorder (ADHD), Allergy (allergic rhinitis), Anemia, Anal fissures, Anxiety, Asthma, BMI, Depression, Endometriosis, Erectile Dysfunction, Fatty liver disease, Fibromyalgia, Food sensitivities, Frailty, Hemorrhoids, High Visceral Adipose tissue (VAT) mass, Hyperlipidemia, Hypertension, IBS, Inflammation metabolic syndrome, Kidney function, Menopausal symptoms, Migraine, Non-melanoma skin cancers, Oral microbiome profile, Osteoarthritis, Osteoporosis, Patterned baldness in men, Polycystic ovary syndrome, Diabetes, Recurrent UTI, Sarcopenia, Skin function, Sleep apnea, Sleep quality, Stress, and Urinary tract stones.

Criteria based on available data for a set of specific features were predefined, representing each health indication, resulting in a total of 202 features. The full list of features within each health indication category, along with the criteria and number of participants available is described in **Supplementary Table 1.**

The health indication characterization used a combination of self-reported conditions, medical diagnostic devices, medical monitoring devices, and medication usage reporting. Below is a short description of the selected health indications and the data used for their definition.

#### Attention deficit hyperactivity disorder (ADHD)

ADHD was determined through self-reporting of a known diagnosis and medication use, specifically ADHD treatments like stimulant medications or centrally acting sympathomimetics. The Eysenck Personality Inventory’s Neuroticism Scale, based on a 12-question survey calculated according to [25], was used as a secondary feature due to some overlaps between neuroticism and ADHD though they stem from different psychological frameworks.

#### Allergy (allergic rhinitis)

Allergic rhinitis was identified through self-reporting in questionnaires and by tracking the use of related medications and associated conditions.

#### Anemia

Anemia was identified through self-reporting and blood test results. Hemoglobin levels below 11.9 g/dL in females and 13.6 g/dL in males, or hematocrit below 35% in females and 40% in males, indicated anemia [26]. Supporting evidence includes taking anemia supplements such as iron, B12, folic acid, and antianemic medications.

#### Anal fissures

Anal-fissures were defined through self-reported diagnosis and recall of fissure surgery, and treatment with specific ointments and muscle relaxants. Additionally, the HPP tracks cases involving frequent hard stools over the past three months, which may be a contributing factor to fissures.

#### Anxiety disorders

Anxiety indication was determined through self-reporting, medication use, and related questions, such as whether a general practitioner or psychiatrist has been consulted for anxiety or related conditions.

#### Asthma

Asthma was determined through self-reported data. Additionally, the presence of asthma was confirmed if participants report taking asthma medications. Supporting data included elevated blood eosinophil counts. History of smoking in the setting of asthma was also assessed to help differentiate asthma from chronic obstructive pulmonary disease (COPD).

#### BMI

Overweight is defined as a BMI of 25 to 29.9 kg/m², while obesity is categorized as a BMI of 30 kg/m² or higher, with severe obesity identified as a BMI of 40 kg/m² or more, or 35 kg/m² with comorbidities [CDC]. Obesity is determined using both categorical and continuous BMI data. Self-reported data on current or past obesity, along with records of bariatric surgery and GLP-1 receptor agonist treatment for obesity, are also used.

#### Depression

Depressive disorders were defined through various sources, including self-reported depression, antidepressant usage, and validated questionnaire indices from the UK Biobank based lifestyle questionnaire, including probable recurrent major depression score [27][PMID: 24282498].

#### Endometriosis

Endometriosis was determined through self-reported diagnoses and surgery history for endometriosis, and menopausal status.

#### Erectile dysfunction

Erectile dysfunction was defined based on self-reporting or specific medications focusing on PDE5 inhibitor drugs like Sildenafil and Tadalafil. Surgical treatments for erectile dysfunction are not asked specifically.

#### Fatty Liver Disease

Metabolic Dysfunction-Associated Fatty Liver Disease (MAFLD) was identified through self-reported diagnoses and confirmed by liver ultrasound assessments that measure the speed of sound through liver tissue, with slower speeds indicating higher fat content. Blood tests measuring liver enzymes like ALT and AST, alongside non-invasive liver fibrosis indices like the Fibrosis-4 (FIB-4) Index, are also used to support the diagnosis.

#### Fibromyalgia

Fibromyalgia was identified through self-reporting in questionnaires and interviews. Additionally, supporting evidence for fibromyalgia included pharmacological treatments, such as tricyclic antidepressants and gabapentinoids, and reported sleep problems and fatigue.

#### Food sensitivities

R Here, food sensitivities were determined through self-reporting food intolerances, allergies, and sensitivities. The indication includes features of food intolerances, lactose intolerance, celiac disease, and life-threatening allergic reactions, though some common allergens are not included in the interview questions hence a bias towards low counts of such allergies is expected as manifested in the number reported.

#### Frailty

Frailty was assessed by a modified Fried Frailty tool [28] through questions about weight loss, exhaustion, hand grip strength, and walking speed.

#### Hemorrhoids

Hemorrhoids indication was determined based self-reported in questionnaires or follow-up interviews and further supported by reports of hemorrhoid-specific treatments or procedures. Supporting evidence included the percentage of human reads in the gut microbiome sample as a high percentage is expected in hemorrhoid patients due to increased epithelial shedding and inflammation, leading to more human DNA in stool.

#### Abdominal adiposity

The abdominal adiposity indication was defined based on DXA scan features including VAT/SAT ratio, VAT area and waist measurements. As well as ratios like VAT index (VAT/height squared) to adjust for general body metrics and WHR.

#### Hyperlipidemia

Hyperlipidemia features are based on blood test values, self-reporting, and use of specific medications.

#### Hypertension

Hypertension indication is defined based on self-reporting, blood pressure measurements, or chronic medications and classified into categories such as non-hypertensive, high blood pressure without diagnosis, suspected hypertension, and confirmed hypertension. Supporting features include systolic blood pressure (SBP) and diastolic blood pressure measurements (DBP).

#### Irritable bowel syndrome (IBS)

IBS indication is determined through self-reporting in baseline interviews or follow-up questionnaires, and it is further assessed through the Digestive Health Questionnaire (UK Biobank) using Rome IV criteria to diagnose IBS and determine subtypes as described in [29]. The HPP also calculates an IBS Symptom Severity Score (IBS-SSS) based on self-reported pain, bowel habit satisfaction, and overall life interference.

#### Inflammation Metabolic Syndrome

Due to limited data on inflammatory markers in the HPP white blood cell count is used as a broad surrogate of inflammation, and in addition, metabolic syndrome presence is evaluated. Metabolic Syndrome is defined using criteria from the National Cholesterol Education Program (NCEP ATP III)[30] and the International Diabetes Federation (IDF) [31]. According to these criteria, a diagnosis is made when three out of five risk factors are met.

#### Kidney Function - Chronic kidney disease (CKD)

The CKD-EPI formula is used to estimate GFR for the CKD indication. Albuminuria is assessed using the urine albumin-to-creatinine ratio (ACR), with stages classified based on the albumin concentration in urine. Blood urea nitrogen (BUN) levels are also reported in the HPP dataset, and the dataset provides CKD stages according to the KDIGO guidelines [32], using the combination of eGFR and ACR for a comprehensive classification of CKD severity.

#### Menopausal symptoms

Menopause was assessed through self-reported data on menstrual cycles and symptoms. Menopausal stages (premenopause, perimenopause, or postmenopause) are categorized based on changes in menstrual patterns and symptoms, while specific symptoms like hot flashes, vaginal dryness, decreased libido, mood changes, and skin dryness are based on answers to the questionnaires.

#### Migraine

The HPP tracks migraine treatment, although over-the-counter medications like Ibuprofen and Acetaminophen may be underreported since they are not listed as chronic prescription drugs. Within the phenotype features, migraines are classified into four grades based on self-reported severity and treatment types, including the use of CGRP antagonists for severe cases. The categorical features include four categories (1) severe migraine with chronic treatment (2) migraine with antimigraine preparations (3) self-reported migraine, untreated and (4) No reported migraine.

#### Nonmelanoma skin cancers (NMSCs)

NMSCs feature is based on self-reported data on Basal cell carcinoma (BCC) and squamous cell carcinoma (SCC) diagnoses and treatments, such as surgical removal or topical fluorouracil therapy.

#### Oral microbiome

Oral microbiome is assessed by calculating alpha diversity using both the Shannon Effective Number and Simpson Reciprocal Index, and beta diversity using the Bray-Curtis dissimilarity index. The microbiome was sequenced from buccal swabs.

#### Osteoarthritis

Osteoarthritis feature is determined through self-reporting, joint replacement surgeries reporting, and taking medications associated with osteoarthritis; however, the real prevalence may be underestimated due to underreporting of symptoms.

#### Osteoporosis

Diagnosis criteria are different by sex and age. In premenopausal women, osteoporosis is typically diagnosed based on fragility fractures and low BMD (Z-score ≤-2.0). For postmenopausal women, osteoporosis is diagnosed based on a T-score of ≤-2.5 on a DXA scan or the presence of fragility fractures [33]. In men under 50, a diagnosis requires both a low BMD for age (Z-score ≤-2.0) and a history of fragility fractures [33]. For men over 50, osteoporosis is diagnosed with a T-score of ≤-2.5 or a fragility fracture, with the hip being the preferred site for BMD measurement. Osteopenia and osteoporosis features are determined based on self-reporting or results of the DXA scan.

#### Patterned baldness in men

The HPP defines Male pattern hair loss (MPHL) using self-reported balding patterns based on the UK Biobank questionnaire, categorizing individuals into four balding types (No Balding, Frontal, Vertex, or Frontal and Vertex) [34] and use of medications for balding.

#### Polycystic ovary syndrome (PCOS)

PCOS was defined through a self-reported questionnaire, asking participants if they have been diagnosed with the condition. PCOS diagnosis is supported by checking for menstrual irregularity current and past.

#### Pre-Diabetes

Prediabetes feature is defined by self-reporting, fasting glucose, HbA1C% values, and use of antidiabetic agents. Glucose regulation is categorized into normal, prediabetes, and diabetes.

#### Recurrent urinary tract infections (UTIs)

Recurrent UTIs feature is defined by self-reported questionnaires, focusing on the frequency of UTIs and use of prophylactic treatments, with supporting evidence also including cranberry consumption (preventive).

#### Sarcopenia

Sarcopenia feature was determined based on DXA scan lean mass of the limbs, appendicular lean mass index (ALMI) categories based on [35], whole body lean mass and handgrip strength.

#### Skin Function

Included two main dermatologic conditions: acne and numerous moles (>50 moles). Acne and multiple skin moles features are defined through self-reporting and medical questionnaires, with specific features identified for acne diagnosis, treatment, and mole counts.

#### Sleep apnea (obstructive sleep apnea, OSA)

The HPP employs the WatchPAT-300 device to measure sleep apnea severity via indices such as the Apnea/Hypopnea Index (AHI), Respiratory Disturbance Index (RDI), and Oxygen Desaturation Index (ODI). OSA features are defined from the sleep measurements and self-reporting, with conservatively defined OSA as moderate to severe OSA with an AHI or ODI ≥15, and more liberally to include mild cases with AHI or ODI ≥5.

#### Sleep quality

Sleep quality features are assessed using both objective measurements from a home sleep apnea test device, and subjective responses from a UK Biobank based lifestyle questionnaire [36]. Sleep metrics used were the number of wakes, percentage of deep sleep, sleep latency, total sleep time, and wake time after sleep onset, providing a comprehensive view of sleep patterns. Sleep Index [36] is based on questionnaires data, ranging from 0 (worst) to 6 (best), based on responses to questions about snoring, sleepiness, sleep duration, insomnia symptoms, and morning alertness, with higher scores indicating better sleep quality.

#### Stress Response

T The stress response features are based on the WatchPAT 300 device overnight recording and derivation of Heart Rate Variability (HRV) data, providing insights into autonomic regulation during sleep. Additionally, we defined a feature looking into changes in blood pressure and heart rate upon standing to diagnose conditions like orthostatic hypotension and postural tachycardia syndrome. Last, features are also defined by self-reporting of fainting, taking drugs with orthostatic effects like beta blockers, and reports about feeling tense and restless in the past 2 weeks.

#### Urinary tract stones (urolithiasis)

Urolithiasis feature is identified through self-reporting and is supported by documentation of shock wave lithotripsy or evidence of hematuria.

### Association Analysis

The association analysis included data from only one visit per participant: per each health indication the microbiome sample with the most complete feature data from each participant was selected.

Species prevalence was estimated as the proportion of participants with a relative abundance greater than 0.0001. Only species with a prevalence of at least 1% were included. For the association analysis, we used both relative abundance (as a continuous variable) and presence/absence (as a binary variable). Continuous abundance values were log10-transformed, with a pseudocount of 4 added to handle zeros.

We conducted the Shapiro-Wilk test for normality on continuous health indications features, post adjusting for age and sex, to determine the need for log transformation. Log transformation was applied when it showed a closer approximation to the normal distribution, as indicated by a higher Shapiro-Wilk test p-value compared to the original data.

Extreme outliers exceeding 8 standard deviations from the mean were removed from the analysis. For binary and ordinal variables, we ensured a minimum class size of five, merging any smaller classes with the next least severe class. Post-collapsing 8 variables did not have a sufficient number of cases (resulting in no variance) and were excluded from the association analysis, resulting in a total of 194 features.

To identify an association between the health indications (treated as the dependent variable) and the gut-microbiome (treated as the independent variable), we employed both parametric (Ordinary Least Squares (OLS)) and non-parametric (Spearman partial-rank correlation) tests, adjusting for age and sex. Depending on the nature of the dependent variable, we selected the appropriate statistical model: OLS regression for continuous variables, logistic regression for binary variables, and ordered logistic regression for ordinal variables. For continuous features, a robust OLS analysis was also conducted using clipping, where the extreme values in the top and bottom 1% were set to the 99th and 1st percentiles, respectively.

Of note, in some cases of binary and categorical features with small class sizes and low-prevalence species (many zeros), the Generalized Linear Model (GLM) failed to converge—yielding no output or producing unreliable results, such as excessively large coefficients and confidence intervals, and leading to insignificant p-values. We flag instances where GLM convergence fails and recommend relying on the Spearman correlation results in these cases. Additionally, we do not provide standardized effect size estimates for instances where GLM did not converge.

We considered an association statistically significant if it exceeded the threshold for indication-wise multiple comparisons, calculated as 0.05 divided by the total number of species (P = 4.2×10^-5^). An indication was regarded as signal “enriched” if 50 different species surpassed this threshold. Species were considered favorable across multiple conditions if they were associated with at least 10 conditions, each showing a p-value < 1 × 10^-4^, in a direction consistent with favorable phenotypic states.

All analyses were performed using Python, rank correlations were calculated using the package pingouin (v0.5.3).

### Enrichment Analysis

We obtained the genomic annotations of Kyoto Encyclopedia of Genes and Genomes (KEGG) functional pathways [37]for each representative species using Prokaryotic Genome Annotation Pipeline (Prokka) [38] and evolutionary genealogy of genes: Non-supervised Orthologous Groups (eggNOG) [39] tools, as described previously [40]. Gene sets were constructed for each annotation term by aggregating the set of species containing that term. In this process, we lost information about the number of genes contributing to each pathway within a species, retaining only species-level presence or absence per pathway. Gene sets were filtered to retain those present in at least five species but in no more than 95% of the species, removing overly rare or ubiquitous terms.

Enrichment analyses were conducted using both overrepresentation analysis (ORA) and a Gene Set Enrichment Analysis like (GSEA) approach. In the first, enrichment of functional terms was tested among the top-ranked species associated with traits of interest using the hypergeometric test. The species were stratified by direction of effect, as this strategy commonly used to uncover more meaningful biological insights. In the second, species were ranked by association scores, and functional enrichment was assessed along the ranked list using permutation testing (n = 1,000). To select species for ORA, we applied two false discovery rate (FDR) thresholds to the OLS results: 0.01 or 0.05. This allowed us to balance the inclusion of sufficient species for phenotypes with weaker signals while minimizing noise for stronger associations. P-values were adjusted for multiple testing using the FDR-adjusted q-values. Enrichment analyses and visualizations were performed using the Python package GSEApy [41], which provided functions for ORA and preranked GSEA, as well as for plotting enrichment results.

## Results

This study aims to provide a comprehensive and systematic exploration of the associations between the gut microbiome and microbiome-related health conditions, establishing a framework for meaningful comparisons. The Human Phenotype Project (HPP) 10K cohort, a large-scale population study with extensive metagenomic data and detailed phenotypic profiling, offers an ideal foundation for systematically identifying microbiome signatures linked to health outcomes. Key steps in this study include: first, defining and constructing robust health indicators by integrating refined phenotypes; next, performing comprehensive association analyses; and finally, identifying microbiome-relevant signals. The primary objective is to make this dataset publicly available to facilitate future research and support the development of microbiome-based therapeutic strategies.

### Definition and Construction of Health Indications for Microbiome Analysis

We identified 37 health conditions as relevant health indications based on the availability of data within the cohort. Most of the conditions have either well-established or emerging associations with the microbiome. Conditions strongly associated with the microbiome include abdominal adiposity [42,43], BMI [44], type 2 diabetes [45], metabolic syndrome and inflammation [46], hyperlipidemia [47], Metabolic dysfunction-associated fatty liver disease (MAFLD) [48], allergy [49], asthma [50], irritable bowel syndrome (IBS) [51], and oral microbiome diversity [52]. Conditions with emerging evidence include attention-deficit/hyperactivity disorder (ADHD) [53], anxiety [54], chronic kidney disease (CKD) [55], depression [54], polycystic ovary syndrome (PCOS) [56], fibromyalgia [57], frailty [58], hypertension [59], menopause [60], migraine [61], osteoarthritis [62], osteoporosis [63], sarcopenia [64], skin function [65], sleep quality [66], stress [67], food sensitivities [68], recurrent urinary tract infections (UTI) [69], and urinary tract stones[70]. A subset of conditions: anal fissures, hemorrhoids, non-melanoma skin cancer (NMSC), patterned baldness, obstructive sleep apnea (OSA), and erectile dysfunction, currently have minimal or no evidence linking them to the microbiome and are included as exploratory outcomes in this study.

To achieve robust association with these health conditions, we integrated comprehensive molecular and clinical data to develop refined phenotypic features. The refined phenotypes were curated by a clinician, resulting in a detailed set of 198 features representing the selected health indications. **Supplementary Table 1** provides a comprehensive overview of the health indications and their corresponding curated features, including details such as class sizes, classification criteria, thresholds, biomarkers, devices used, and other relevant methodological information. For additional context, please refer to the Methods section. **Supplementary Figure 1A** displays the number of curated features and the distribution of data types (binary, continuous, and ordinal) per each health indication. **Supplementary Figure 1B** shows the number of unique samples available for analysis per each curated feature. Notably, the sample size varies depending on data availability, which is intrinsic to such population cohort studies.

### Association between Health Indications and the Gut Microbiome

At the time of this study the HPP-10K gut microbiome dataset comprised 3,209 gut microbiome species across 10,623 samples obtained from 8,783 participants after quality control procedures (see Methods). In addition to species-level abundance data, we calculated gut microbiome diversity metrics, including Shannon and Simpson alpha diversity indices to quantify species richness and evenness within individual samples, and Bray-Curtis dissimilarity to assess beta diversity, reflecting differences in microbial composition between individuals (**Supplementary Figure 2**). These diversity metrics offer insights into the overall relationship between the gut microbiome and health indicators, complementary information to the changes in individual bacterial taxa. Notably, the diversity metrics showed strong within-person correlation over a two-year period (r ≥ 0.6), as observed in 1,782 participants with a follow-up sample (**Supplementary Figure 2B**). This highlights the temporal stability of the gut microbiome, supporting the use of a single timepoint to represent an individual’s microbiome composition; however, reliance on a single stool sample may still miss short-term fluctuations or transient exposures that influence microbiome composition.

### Individual Species Analysis

Species with at least 1% prevalence were analyzed to ensure sufficient statistical power for association analyses resulting in 1,184 species analyzed. Association analyses were performed using both a binary feature of the presence or absence of species and continuous relative abundance levels. Continuous abundance levels were log-transformed to normalize the distribution. A single visit per participant with both microbiome and phenotypic information was selected, resulting in the number of samples shown in **Supplementary Figure 1B**. Statistical analysis included both parametric and non-parametric tests based on the nature of the variables. Ordinary Least Squares (OLS) regression was used for continuous variables, logistic regression for binary variables, and ordered logistic regression for ordinal variables. Robust OLS analysis was also conducted by clipping extreme values at the top and bottom 1% to the 99th and 1st percentiles. All analyses were adjusted for age and sex. Complete association results for all analytical models are presented in the **Associated Results Folder.**

The statistical significance of the parametric association analyses between individual species and health indications is visualized in an inverted Manhattan plot **(Figure 1)**. Several health indications showed an excess of significant associations beyond what would be expected under the null hypothesis, indicating strong relation between the gut microbiome, defined as more than 50 species surpassing the indication-wise threshold P = 4.2×10-5 (Bonferroni correction for the total number of species analyzed 1,184). Among these, conditions with well-established links to the gut microbiome, such as abdominal adiposity, BMI, hyperlipidemia, metabolic syndrome, MAFLD, IBS, diabetes, and hypertension, displayed pronounced signals.

**Figure 1:**
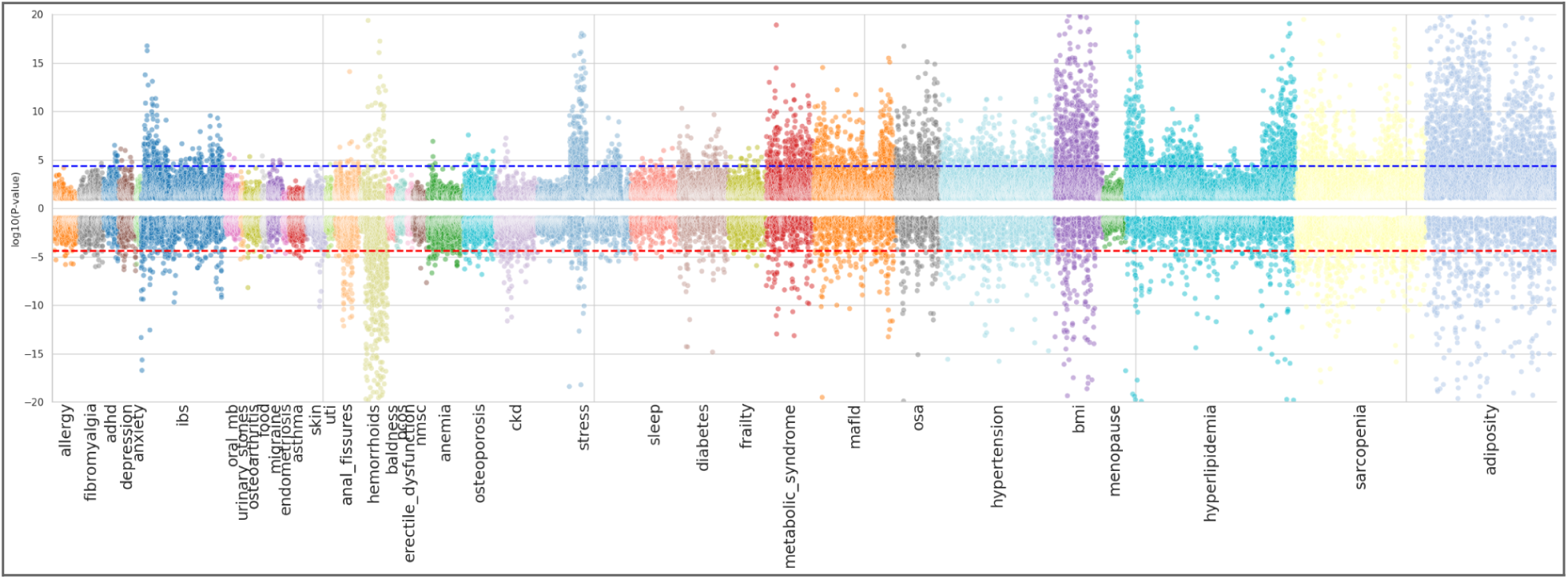
Inverted Manhattan Plot of Microbiome-Health Indication Associations. The inverted Manhattan plot displays the log-transformed p-values for associations between microbiome features and multiple health indications. Each point represents a microbial species, with colors indicating different health indication categories. Dots plotted above the center line (-log₁₀(p-value)) reflect species whose increased abundance is associated with favorable health states, while dots plotted below the center line (log_10_(p-value)) indicate associations with unfavorable health states. The dashed horizontal line indicates the threshold for statistical significance, calculated as 0.05 divided by the total number of species (P = 4.2×10^-5^). The y-axis was clipped to -log10(p-value)= 20 to ensure that all indications are visible.

Additional associations were also evident in conditions with emerging or suggestive associations, including sarcopenia, stress, CKD, and OSA. Hemorrhoids and anal fissures, which have limited prior evidence supporting a microbiome connection, also showed strong associations. However, these associations should be interpreted with caution, as their top features - “percentage of human reads in the microbiome” and “hard stool”, are features that are closely linked to the microbiome sampling process and may reflect confounding rather than a biological association. The strongest feature of each health indication is highlighted in **Supplementary Table 1.** We classified the favorable and unfavorable direction for each feature as shown in **Supplementary Table 1**, while acknowledging that for some features this distinction is not straightforward, as the interpretation may vary depending on the clinical or physiological context.

### Diversity Metrics Analysis

The associations between the gut microbiome diversity metrics and the health indications are presented in an inverted Manhattan plot for easy comparison to the species-level analysis in **Figure 2**. We observed that the associations with alpha diversity (species richness and evenness) closely align with the enrichment observed in the species-level association analysis, with strong signals observed for the same health indications (except for Anal Fissures). Significant associations with alpha diversity at P-value < 4.2×10^-5^ were found with: Abdominal adiposity, Hemorrhoids, Sarcopenia, BMI, Hyperlipidemia, Stress, Hypertension, MAFLD, IBS, Metabolic Syndrome, OSA, Diabetes, and CKD, with the addition of ADHD being borderline significant. Notably, only three indications: Hemorrhoids, BMI, and Hypertension, showed a significant association with beta diversity at the threshold for P-value < 4.2×10^-5^.

**Figure 2:**
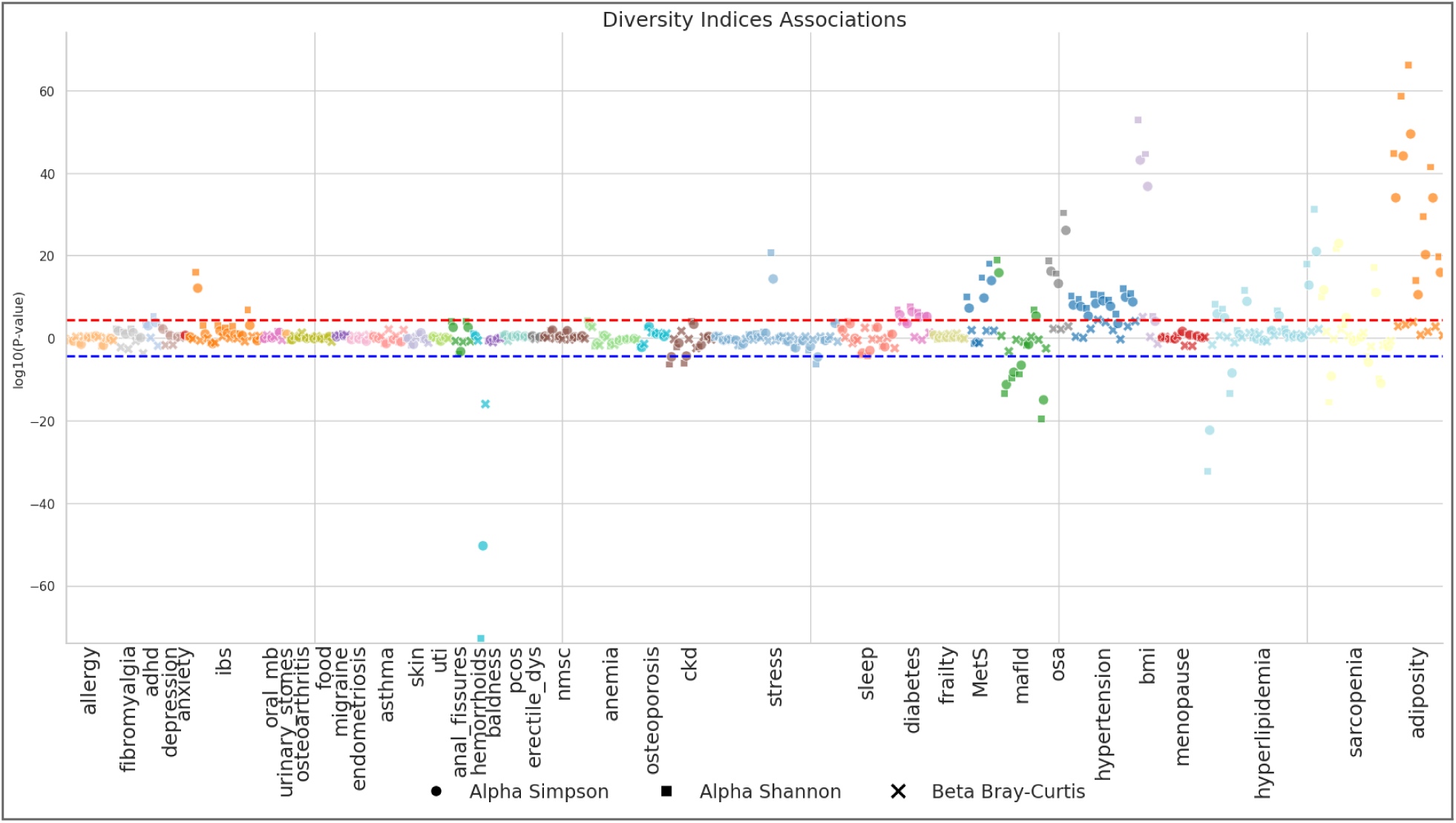
Diversity Indices Associations with Health Indications. The inverted Manhattan plot displays the log-transformed p-values for associations between gut microbiome diversity indices and health indications. Dots plotted above the center line (-log₁₀(p-value)) indicate that diversity increases with favorable health states, while dots plotted below the center line (log_10_(p-value)) indicate that diversity increases with unfavorable health states. Each shape denotes a different diversity measure: Shannon diversity (X), Simpson diversity (O), and Bray-Curtis beta diversity (□).

The direction of effect for the association with alpha diversity indicated higher diversity values in favorable condition states, suggesting that greater microbial richness and evenness are associated with more favorable health profiles. Similarly, at the species level analysis, many more species showed significantly higher abundance in favorable states compared to those associated with unfavorable states. This pattern is evident in **Figure 1** and in the supplementary volcano plots (**Associated Result Folder**). Together, these associations provide complementary evidence linking gut microbiome composition to these health indications.

### Enrichment Analysis

Enrichment analysis is a commonly used method for exploring large-scale association results and it highlights the types of downstream analyses enabled by this dataset. To explore whether the species-level signals clustered into functionally meaningful pathways, we performed KEGG pathway enrichment analysis. This approach serves both to validate the microbiome-health indication associations and provide insight to further mechanistic interpretation. Species were assigned to 229 KEGG pathways at the gene level using Prokka followed by EggNOG annotation [38,39]. Any species containing at least one gene assigned to a KEGG pathway was considered part of that pathway. Thus, our approach does not account for the number of pathway-related genes within a species, meaning that a species with a single pathway-related gene is weighted equally to one with many such genes. We employed over-representative analysis (ORA) stratified by direction of effect. **Figure 3** illustrates examples of KEGG pathway enrichment results plots; full results across all indications are provided in the **Supplementary Table 2.** The Dot-plot in **panel A** shows enriched pathways (FDR < 0.05) among species associated with favorable abdominal adiposity state, based on a species selection threshold of adjusted p-value < 0.01. This threshold resulted in hundreds of species, enabling ORA. The Bar-plot in **panel C** shows enriched pathways for species associated with both favorable and unfavorable frailty states. For this indication, the adjusted p-value < 0.01 threshold was too stringent and yielded no species; therefore, we relaxed the threshold to 0.05, identifying approximately 50 species for ORA. To address such limitations of threshold optimization, we also show gene set enrichment analysis (GSEA), a method that evaluates functional enrichment across the entire ranked species list without requiring a threshold. **Panel B** illustrates GSEA results for three enriched pathways associated with abdominal adiposity. Notably as observed, certain pathways, such as ‘bladder cancer’, appear across multiple traits. Pathways that are enriched across most indications probably represent artifacts in the species-set definitions and should be interpreted cautiously or excluded from downstream interpretation.

**Figure 3:**
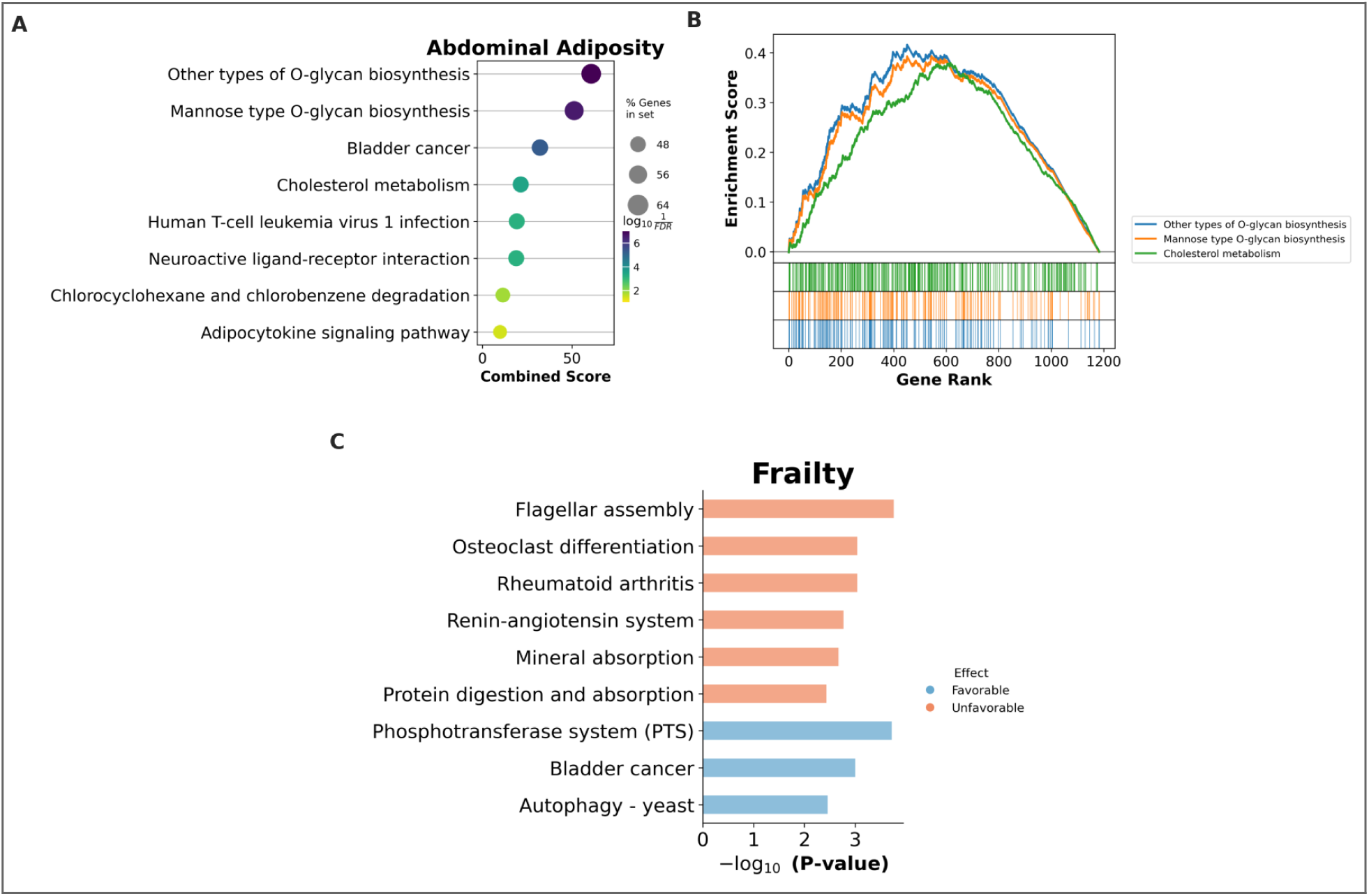
Pathway enrichment of microbiome species associated with health indications. A. Overrepresentation analysis (ORA) results for Abdominal Adiposity, showing KEGG pathways enriched among species favorably associated with the trait (adjusted p-value < 0.01). The dot plot displays the top pathways (FDR < 0.05), where dot size indicates the percentage of genes in each pathway and color reflects enrichment significance (−log_10_FDR). B. Gene Set Enrichment Analysis (GSEA)-like enrichment curves for three KEGG pathways enriched in Abdominal Adiposity. Species were ranked by Spearman correlation coefficient, and enrichment scores were computed across the full ranked list without the need for a threshold. Tick marks below each curve indicate pathway-member species positions. C. ORA results for Frailty, based on species associated at an adjusted p-value < 0.05. The bar plot presents enriched KEGG pathways stratified by direction of effect. Salmon bars indicate pathways enriched among species more abundant in higher frailty (Unfavorable state), while blue bars indicate enrichment among species more abundant in lower frailty (Favorable state).

### Core Species Associated Across Health Indications

We aimed to identify key microbial species consistently associated with favorable health states across multiple indications. Cross-indication evaluation pinpointed several species significantly linked at P-value < 1 × 10-4 to 10 or more distinct health indications, highlighting these as potential core or foundational taxa. **Figure 4** illustrates species consistently associated with favorable health states (n=57), alongside species (n=10) linked to unfavorable states to help guide efforts that may aim to promote or deplete specific taxa. For each indication, we show the Spearman correlation with the most strongly associated feature within that indication. Associations with favorable states are shown in blue, and those with unfavorable states in red. Note that for some features, classifying states as favorable or unfavorable is not always straightforward, as this may vary depending on the context.

**Figure 4:**
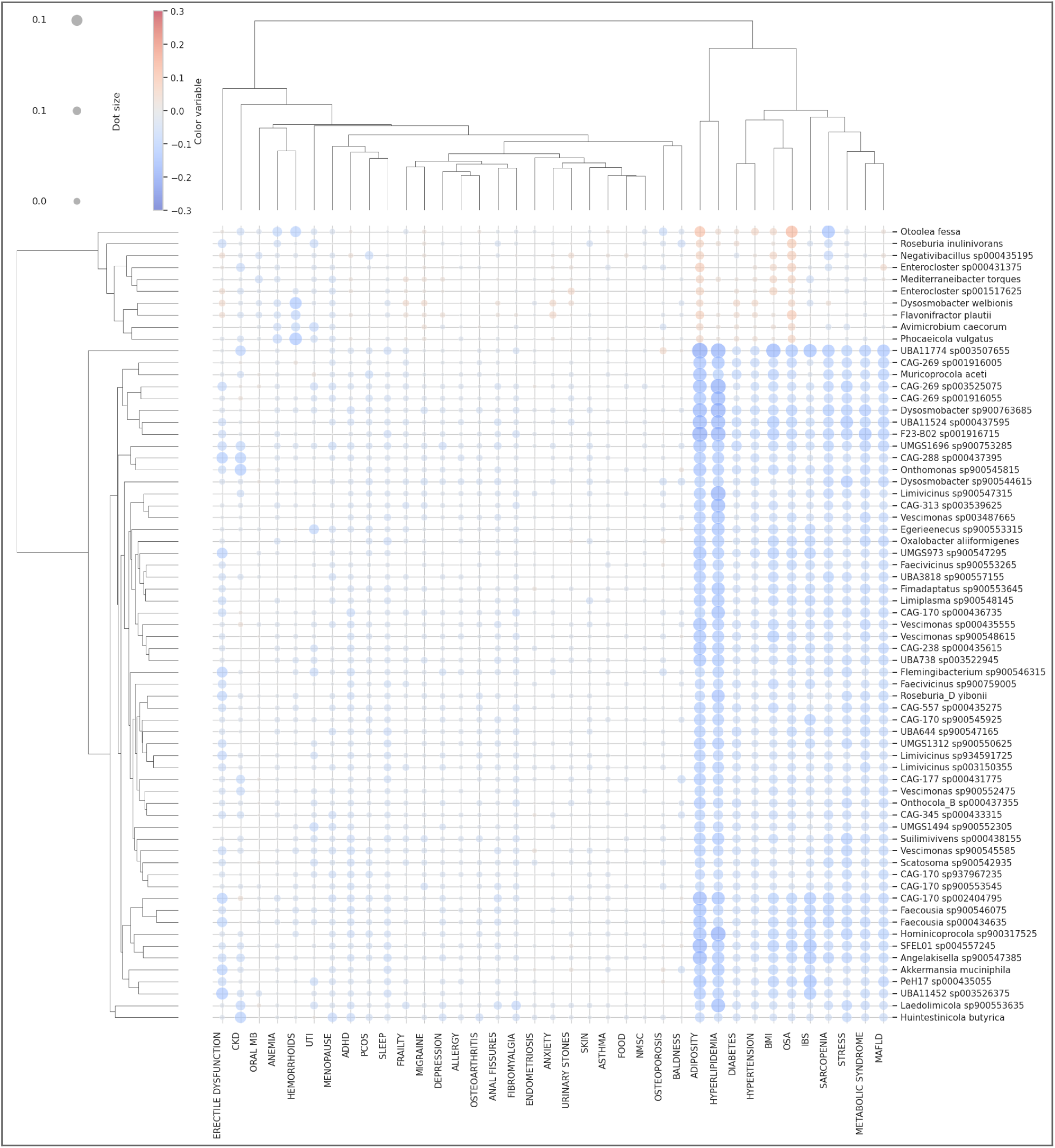
Species with Significant Associations Across Multiple Health Indications. Dot plot of Spearman rank correlation coefficients between microbial species and health indications for species with significant (P-value < 1 x 10 ^-4^) associations across multiple indications (≥10 indications). Each row represents a species and each column an indication. Dot size reflects the strength of the correlation, and dot colors indicate the direction of association: blue for associations with favorable health states, red for associations with unfavorable health states. Absence of a dot indicates no substantial correlation (|r| < 0.001). The correlation presented corresponds to the most significant feature for each indication. The species at the top of the plot are consistently associated with unfavorable health states (n = 9), whereas those at the bottom are consistently associated with favorable health states (n = 42). Correlations are shown for the most significantly associated feature within each indication.

To illustrate the utility of our comprehensive dataset, we present five species that show consistent associations with favorable health states at the feature level (**Figure 5**). We focused on species from taxa with existing scientific literature to assess whether our data can offer new insights. *Blautia* is one of the major intestinal bacteria, a member of the Lachnospiraceae family, and is known to produce butyrate, a short-chain fatty acid that supports intestinal homeostasis, energy metabolism, and has anti-inflammatory properties [71]. It has drawn attention for its potential influence on metabolic processes and fat storage, particularly visceral fat [72,73]. In line with these roles, our data identify *Blautia sp900066205* as our most strongly associated *Blautia* species, showing consistent correlations with lower VAT, triglyceride levels, and BMI. *Oscillospiraceae CAG-170 sp002404795* and *Vescimonas sp000435555*, from a less well-known genus, were identified as core species consistently associated with favorable health states. They are members of the class Clostridia. Prior research has reported a negative correlation between the abundance of the CAG-170 and Vescimonas genus and BMI [74]. Both species show strong favorable correlations with VAT, triglyceride levels, and additional metabolic features at the species level. These findings reinforce previously observed genus-level associations and highlight specific species.

**Figure 5.**
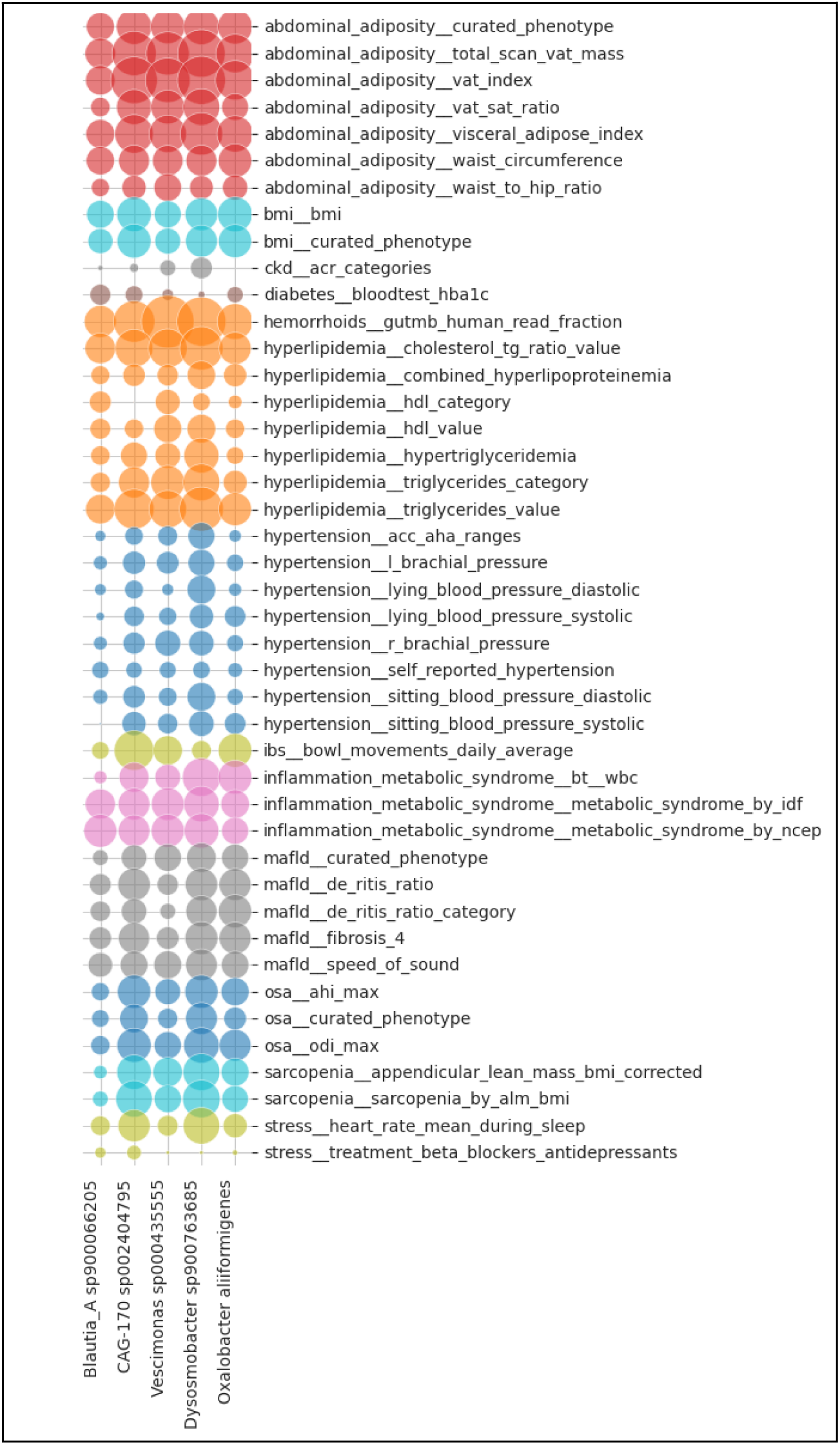
Feature-level associations of five literature-supported species associated with favorable health states. Dot plot showing Spearman correlation coefficients between five microbial species and selected health features. Each column represents a species, and each row corresponds to a health feature. Dot size indicates the strength of the correlation, while color represents the health indication. All correlations are in the favorable direction. Only features with multiple significant associations are displayed to improve clarity given the large number of features. These species were chosen to illustrate the utility of our dataset for high-resolution exploration of species–health associations supported by prior literature.

The genus Oxalobacter has been reported with both beneficial and harmful associations related to cardiometabolic conditions, such as T2DM and CAD. While some studies suggest that increased Oxalobacter abundance may elevate the risk of T2DM and CAD [75, 76], others have reported a potential protective role, particularly in connection with reduced vascular calcification and improved metabolic outcomes following bariatric surgery [77, 78]. In our analysis, we observed a consistent favorable correlation specifically with the species *Oxalobacter aliiformigenes* across multiple health indications, including diabetes and cardiovascular-related traits such as Hypertension, Hypercholesterolemia and HbA1c. This species-level resolution may help reconcile previously conflicting genus-level findings.

The species *Dysosmobacter sp900763685* showed favorable correlations across multiple health indications, most notably with VAT and triglyceride levels. It belongs to the same genus as *Dysosmobacter welbionis,* which has been described in the literature as supportive of metabolic health (particularly in mice) [79]. However, in our analysis, Dysosmobacter welbionis was significantly associated with unfavorable states across several metabolic features. These findings are noteworthy, as this species is suggested as a candidate for ‘next-generation probiotic’ [80].

Among other species identified in our analysis as being associated with multiple unfavorable metabolic states (as shown in Figure 4), *Mediterraneibacter torques* (formerly Ruminococcus torques) and *Negativibacillus sp000435195* have both been previously implicated as harmful in metabolic and inflammatory conditions. *Mediterraneibacter torques* have been linked to disorders such as inflammatory bowel disease and obesity [81], and *Negativibacillus sp000435195* has been associated with fatty liver disease [82]. Our findings further reinforce these adverse associations. Moreover, in the case of *Phocaeicola vulgatus*, the literature presents mixed results [83,84]; whereas *Enterocloster sp000431375* was previously considered potentially beneficial in the context of fatty liver disease (referred to as Clostridium M sp000431375) [82]. In our data both showed associations with several adverse metabolic states, particularly those related to adiposity. These examples highlight the value of systematically querying a wide range of microbial species across diverse health outcomes in a healthy population,offering broad insights into microbes of potential interest for future investigation.

## Discussion

We investigated the relationship between the gut microbiome and a wide range of health indications, aiming to create a comprehensive and systematic resource for uncovering microbiome-related associations. To achieve this, we selected 37 microbiome-relevant health indications and integrated detailed molecular and clinical data to define refined phenotypic features, resulting in a curated set of approximately 200 traits. The use of the deeply phenotyped and extensively sampled HPP cohort was critical, as precise phenotype definition is essential for identifying robust and biologically meaningful associations. This dataset represents a uniquely rich resource, offering an opportunity to explore microbial links to human health and guide the development of targeted, evidence-based probiotic interventions.

The ability to systematically assess a wide range of health indications and measurement types within a single, deeply phenotyped cohort is a key strength of this study. This uniformity reduces the potential biases that often arise when comparing across separate studies with heterogeneous populations, protocols, or sequencing platforms. By analyzing multiple conditions under the same study design, microbiome processing pipeline, and statistical framework, we enable more reliable cross-condition comparisons. This integrated approach offers a robust foundation for identifying core microbiome features relevant across different health domains and supports the utility of the dataset as a broad resource for future investigations.

We observed a significant enrichment in associations between the gut microbiome and various health indications (n = 14). An indication was considered enriched if at least 50 microbial species surpassed the significance threshold for multiple species comparisons (P = 4.2×10^-5^). This threshold, is somewhat permissive, as the study is exploratory, and intended to serve as a resource for identifying promising indications and species for more detailed and rigorous future investigations. Notably, due to the inherent correlation structure among microbial species and phenotypic features, defining a strict universal threshold is challenging. Importantly, the majority of the associations were of favorable relationships, with species being more prevalent in individuals with favorable phenotypes compared to those with unfavorable. Given that the HPP is a relatively healthy population cohort, it may offer a unique advantage in identifying potentially favorable species that could be targeted for probiotic development.

While the most pronounced conditions exhibited the strongest associations, other indications may still contain meaningful microbiome relationships. For 23 indications, no prominent associations appeared in the Manhattan plot, potentially due to small sample sizes, less informative phenotypes, or low condition prevalence. Nevertheless, some of these indications showed consistent microbial signals across both continuous and binary abundance analyses and may benefit from follow-up as the HPP cohort continues to grow and follow participants longitudinally.

A key limitation of this study is the variability in sample sizes across conditions, which is determined by the availability of data within the cohort. This variability should be considered when interpreting the strength of associations, as smaller sample sizes may reduce statistical power and lead to less robust findings. Furthermore, the number of features per condition and the frequency of conditions vary. Conditions with fewer features or lower prevalence may give rise to less comprehensive or generalizable results, making it difficult to directly compare associations across conditions. However, these challenges are inherent to population-based cohorts, which, by design, lack selection criteria representing real-world variability. Also, The relatively healthy and health-conscious nature of the cohort may introduce a healthy volunteer bias, potentially underestimating associations with more severe disease phenotypes. Despite these limitations, the use of a large population cohort with deep phenotypic profiling remains a comprehensive approach for investigating microbiome-related health associations.

Another limitation inherent to microbiome data is its compositional nature, which can introduce biases in analysis and interpretation. To mitigate this, we employed multiple analytical approaches, including both continuous species abundance and presence-absence analyses, using parametric and non-parametric approaches. We believe that true biological signals should remain consistent across analytical approaches and recommend that researchers using this dataset apply this cross-validation method. Finally, we note the potential subjectivity involved in defining the favorable direction of each health feature. While a clinician curated the directionality based on standard clinical interpretations, some phenotypic features may not have a universally agreed-upon favorable or unfavorable state, and interpretations can vary depending on the specific clinical context.

Large-scale, high-throughput microbiome studies, such as ours, deal with the challenge of how to effectively aggregate and interpret the extensive volume of results. We illustrated the use of species set enrichment analysis as one strategy to add functional and biological context to species-level associations. While this approach can highlight biologically meaningful patterns, it also produces a high rate of false positives, due to the nature of hypergeometric testing across large numbers of species sets or the limitations in the precision and reliability of the species sets. Our species sets were derived from genome-level annotations, using Prokka and eggNOG to assign orthologous KEGG pathway annotations. This method aggregates gene-level data to species level; losing quantitative gene abundance information, which could introduce noise. To circumvent one approach is to perform pathway analysis directly at the gene level, as we have done in our previous work [85]. However, in the present study, we chose to maintain species-level annotation to align with the focus on microbial taxa, the species sets are available as a resource, and researchers are encouraged to build upon or refine them to suit their specific research questions.

We also explored the data by species to identify microbes consistently associated with favorable health indications across multiple conditions. This analysis revealed several species linked to favorable phenotypic profiles, suggesting they may play a meaningful role in supporting overall health and could be promising candidates for future research and therapeutic development. However, it’s important to consider the phenotypic correlations between different health features when interpreting the results of these species to better understand the potential health benefits. In particular, the association with strong interrelated features such as VAT can add a layer of complexity. VAT is often correlated with a broad range of health outcomes and can act as a confounder. Accounting for these effects is challenging but essential for improving our understanding of causal relationships and better highlighting potential microbial contributors to health.

To exemplify the potential research applications of our findings, we reviewed existing literature on species consistently associated with favorable or unfavorable health states, noting that relatively few species have been consistently linked to unfavorable states. Our research provides species-level resolution for genera previously implicated in health associations, potentially enabling the characterization of specific taxa that contribute to these associations, such as *Blautia sp900066205, Oscillospiraceae CAG-170 sp002404795* and *Vescimonas sp000435555*. Moreover, our results challenge prior findings such as for *Dysosmobacter welbionis*, highlighting the value of comprehensive population-level analysis in refining or reassessing microbial health associations.

To conclude, this study investigated the relationship between the gut microbiome and a wide range of phenotypic features spanning 37 health indications, with the goal of creating a publicly available resource to support the broader scientific community in identifying promising targets for probiotic development. We found significant species-level associations in 14 of these health indications. Notably, several species favorably correlated with multiple health indications, emphasizing their potential as fundamental species for use in various formulations or in promoting overall health. As this study is observational, associations identified should not be interpreted as causal. Further experimental or interventional studies will be required to determine mechanisms. The next step will be to validate these associated phenotypes and conduct a more detailed analysis to confirm which bacteria are beneficial and to further assess their potential for developing effective probiotic interventions.

## Acknowledgments

We thank Dirk Gevers for providing strategic advice at the inception of this project and for playing an active role in data analysis and manuscript preparation. We are also grateful to Ayya Keshet for her valuable assistance in identifying references and supporting the literature search. We thank all Pheno.AI data science members for useful discussions.

## Ethical approval

All participants signed an informed consent form upon arrival to the research site. All identifying details of the participants were removed prior to the computational analysis. The 10K cohort study is conducted according to the principles of the Declaration of Helsinki and was approved by the Institutional Review Board (IRB) of the Weizmann Institute of Science.

## Data availability

The individual user data used in this paper is part of the Human Phenotype Project (HPP) and is accessible to researchers from universities and other research institutions at: https://humanphenotypeproject.org/data-access. Interested bona fide researchers should contact to obtain instructions for accessing the data.

## Author contribution

D.W-V. led the project planning, data analysis, interpretation of results, and manuscript writing. Y.R. led the derivation of clinical variables and contributed to project planning, result interpretation, manuscript writing, and critical review. T.S. contributed to project planning and was responsible for developing and coding clinical variables. A.J. and A.D. assisted with the implementation and coding of clinical variables. A.G contributed to data acquisition and validation. R.D. conceptualized the project. E.S. conceived and supervised the project, and contributed to study design, interpretation of results, manuscript writing, and critical review.

---

**Supplementary Figure 1:**
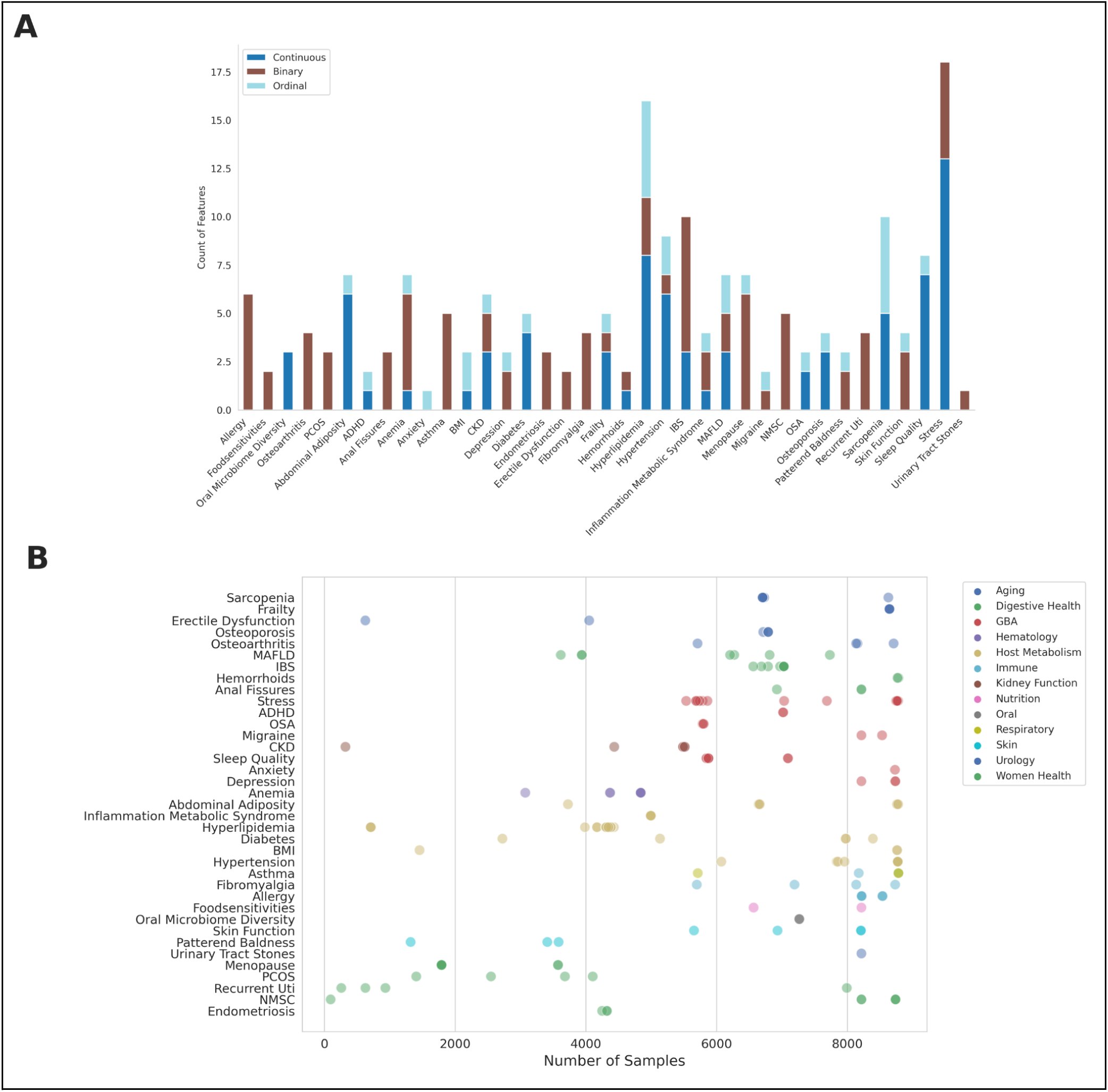
Curated Features per Health Indication. <u>A. Number of Curated Features per Health Indication by Data Type</u> Bar charts display the number of curated features associated with each health indication, categorized by their data type: Binary, Continuous, and Ordinal. Bars are stacked to represent the total count of curated features of a given health indication. <u>B. Number of Unique Samples per Curated Features by Health Indication and Category</u> Each dot shows the number of samples included in the association analysis of a specific curated feature by health indication (row), health indications are color-coded by category.

**Supplementary Figure 2.**
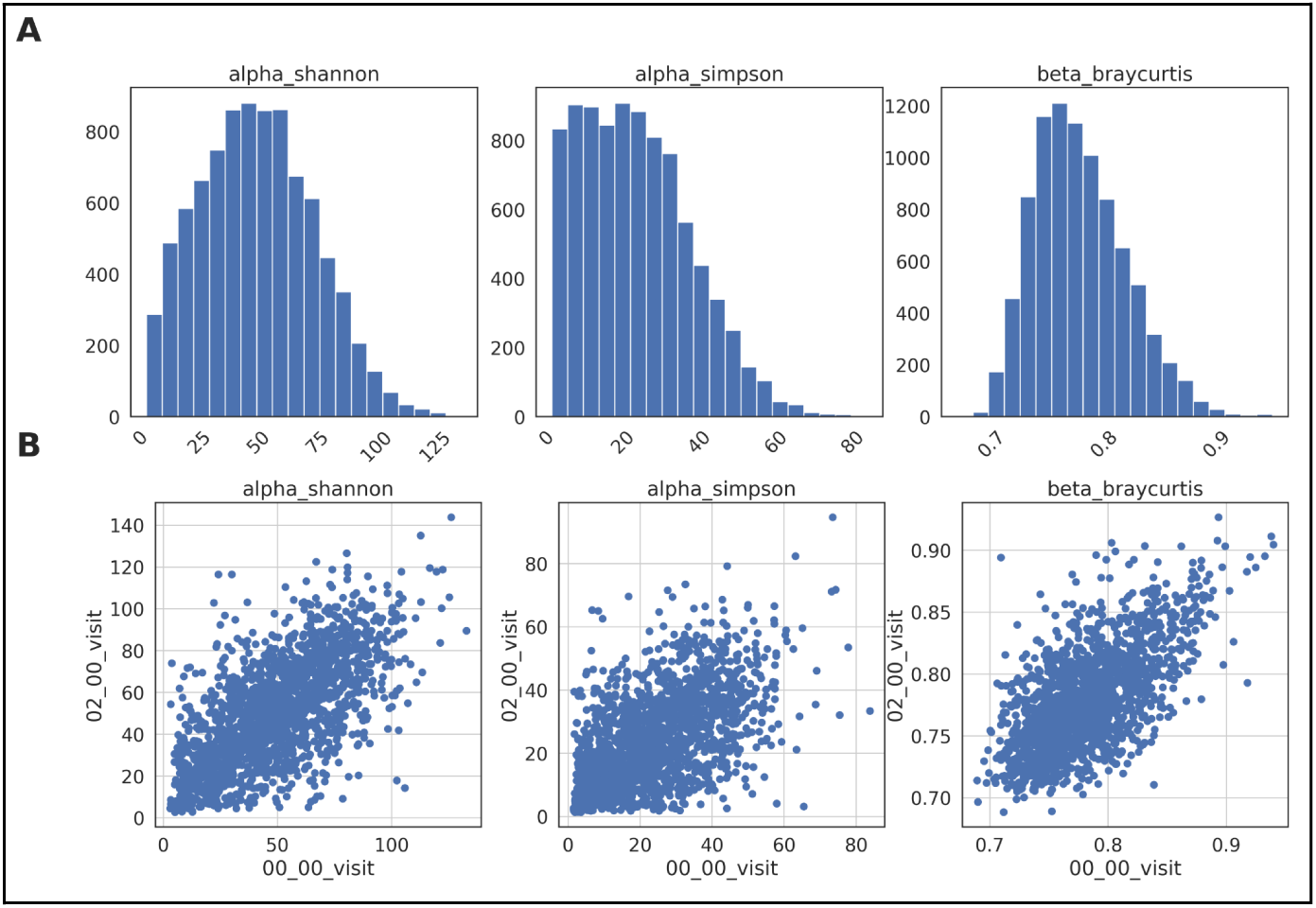
Distribution and Within-Person Correlation of Microbiome Diversity Metrics. A. Histogram of the frequency distribution of gut microbiome diversity metrics in the HPP-10K study population.B. Scatter plots of the within-person correlation of diversity metrics across two-year visits. Each point represents an individual, with the x-axis indicating the first visit (00_00_visit) and the y-axis indicating the corresponding diversity feature at the second visit (02_00_visit). These plots highlight the stability of individual diversity metrics over time.

